# Nonlinear Stability Analysis for Artificial Kidney Multi-compartmental Models

**DOI:** 10.1101/2020.11.26.400606

**Authors:** Rammah Abohtyra

## Abstract

This paper addresses a new global stability analysis for a specific class of nonlinear multi-compartment models with non-positive flows and whose balances are described by equilibrium sets. We apply the stability analysis to our physiological-based model of extracellular fluid used during dialysis therapy in end-stage kidney disease patients. To gain an in-depth understanding of the risk associated with fluid removal by the artificial kidney during the short time (3-5hrs) of the dialysis therapy, we use the stability results to analyze the solution’s behavior of our model under standard ultrafiltration and patient-specific ultrafiltration profiles. Therefore, the standard ultrafiltration profiles do not guarantee optimal outcomes, and we highly recommend incorporating physiological insights into the ultrafiltration profiles to improve outcomes.

## 1. Introduction

Patient in end-stage kidney disease are often suffering from multiple acid-base derangements and fluid shift complications [1, 2]. Dialysis is a life-sustaining treatment that corrects acid-base derangement and removes fluid accumulation to obtain dry weight (normal weight without extra fluid) [3, 4, 5, 6]. During dialysis treatment, an artificial kidney is used to remove excess fluid (2-5 liters) from the blood (plasma) by ultrafiltration, within 2-5 hrs, leading to fluid refilling from the interstitial compartment to overcome the fluid shift [7, 8]. An inadequate balance between fluid removal and refill is associated with increased morbidity and mortality risk [9].

The standard ultrafiltration protocol applies high constant rates, generating fluid imbalance between fluid removal and refill that causes a significant reduction in the blood volume, especially at the end of the therapy when the vascular refilling is low [10, 11]. This fluid imbalance provokes intradialytic hypotension (IDH), (intradialytic systolic blood pressure (SBP) *<* 90 mmHg) in 25-60% of hemodialysis treatments, and is associated with a higher risk of cardiovascular morbidity and rate of death (20-25% death rate each year under hemodialysis maintenance therapy) [12, 11].

Stability analysis, widely used in system biology and control systems, is an essential tool for analyzing complex mathematical models, but it is overlooked in extracellular fluid models whose fluid exchanges are defined by non-positive flows and whose stability is described by equilibrium sets. The current emphasis on improving the assessment of dry weight and strategies for fluid removal in dialysis therapy motivates the need for exploring fluid management and equilibrium to provide possible stabilizing procedures that may offer guidelines for optimal fluid removal strategies in end-stage kidney disease patients.

The total body fluid is distributed mainly between two compartments, the extracellular compartment (ECC), which contains fluid outside the cells including interstitial fluid and plasma (about 20% of the body weight), and the intracellular compartment (ICC), which contains fluid inside the cells, (about 40% of the body weight) [13]. The ECC is divided by a capillary membrane into the interstitial compartment and the intravascular compartment. The intravascular compartment contains blood, which consists of cells (e.g., red blood cells) and plasma. The plasma is the noncellular part of the blood [14, 15]. Fluid-exchange between the intravascular and interstitial compartments is controlled by the Starling forces; namely, the osmotic and hydrostatic pressure gradients [16, 17]. If the difference between these gradients is negative, there will be a fluid filtration across the capillary membrane into the intravascular compartment. However, if the difference is positive, resulting in the filtration of fluid into the interstitial spaces.

Over the past years, there has been an active research effort to characterize fluid removal from the intravascular compartment by ultrafiltration and vascular refilling from the interstitial spaces during dialysis treatment. Two-compartment models describing the short-term dynamics of vascular refilling and ultrafiltration have been derived in (e.g.,[11, 16, 18, 19, 20]). For example, the model in [16] provides a simplified linear two-compartment model and nonlinear two-compartment model incorporating the lymphatic system. The nonlinear model addresses the contribution of lymph flow to the fluid movement from interstitial spaces to the intravascular compartment. More recently, in [11], a nonlinear two-compartment model was developed to describe the vascular refilling during dialysis. This model considers (microvascular) fluid shifts between the compartments and lymph flow from the interstitial to the intravascular compartment.

There are many methods for analyzing the stability of nonlinear compartmental models [21] and others. Among them is the Lyapunov stability approach. Exploring the stability of the nonlinear compartment model with the Lyapunov method is a challenging task because it often requires searching for Lyapunov candidates by trial-and-error. The stability of a class of nonlinear compartment models, based on the Lyapunov approach, has been investigated in [22, 23, 21]. Since the fluid flow of our model is non-positive, the results of these papers cannot be applied to our model because of the particular class of models under consideration.

A class of nonlinear compartment model, similar to our model, has been discussed in [24, 25]. Sufficient conditions concerning the flows between the compartments are made (positive flow). These results, however, do not seem to be applicable to our model because the conditions are not satisfied, and the Lyapunov functions used are inappropriate for our model. In addition, another class of nonlinear models has been studied in [26]. Although these models have a similar structure to ours, the result of [26] is not applicable here because it pertains only to isolated equilibrium points. The above compartmental models were used with external inputs described by a specific class of constant non-decreasing input and with sufficient conditions to extend stability results.

Our contribution in this paper is two folds. First, we overcome these limitations and provide a new Lyapunov function to prove the global stability of a specific class of nonlinear compartment model whose flow functions are non-positive. These results are applied to our two-compartment model used for dialysis therapy in patients with kidney failure. Second, to gain a physiological insight into the risk associated with fluid removal by the artificial kidney, we extend the results to include the effect of the input (ultrafiltration) on the model to investigate the boundedness and behavior of the solution of the nonlinear model during ultrafiltration.

We organize the paper as follows: In Section 2, a brief description of an artificial kidney model is presented. In Section 4, a global stability analysis is provided, including the main results. In Section 5, simulation examples are provided. Finally, in Section 6 is the conclusions.

## 2. Artificial Kidney Model description

The two-compartment model represented by Fig. 1 is comprised of the intravascular and interstitial compartments, which are separated by a capillary membrane wall, and lymphatic flow, which returns a small amount of fluid from the interstitial compartment into the intravascular compartment [27]. Because the plasma is the extracellular fluid part of the blood in the intravascular compartment, in this paper, we consider that the plasma volume is part of the intravascular compartment volume. There is a continuous microvascular fluid filtration/refilling between the intravascular and interstitial compartments via the capillary membrane where *K*_*f*_ refers to filtration/refilling coefficient [L/min.mmHg].

**Figure 1:**
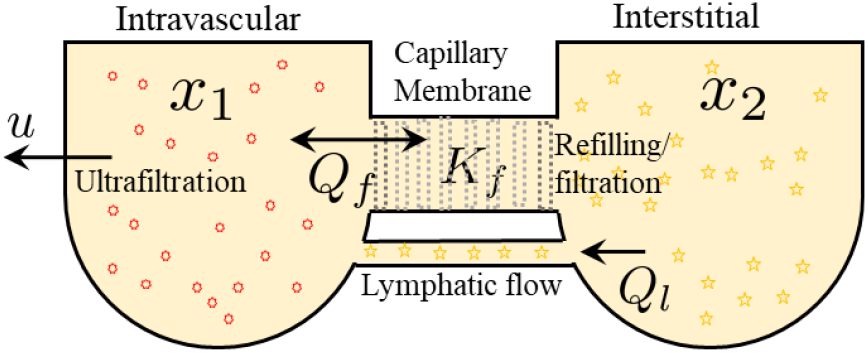
Two-compartment model used for the artificial kidney treatment, contains: interstitial and intravascular compartments separated by a capillary membrane, and includes lymph flow and ultrafiltration.

- Input: The ultrafiltration rate (UFR) is the external input to this model (denoted by *u* with negative sign), which describes the fluid removal from the intravascular compartment (i.e., plasma) during dialysis treatment.
- Output: The hematocrit (*Hct*) is the measured output, which is the proportion of the total blood volume that consists of red blood cells (RBCs); *Hct* is usually expressed as a percentage in clinical practices but is defined here as a fraction.

The mathematical description of this model, during dialysis, is given by nonlinear differential system:

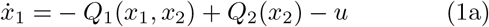

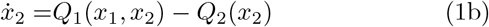

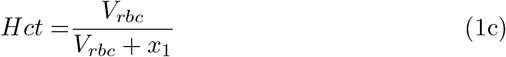

where *x* = (*x*_1_, *x*_2_) is the state of the model, *x*_1_ and *x*_2_ represent the intravascular and interstitial volume, respectively; and *V*_*rbc*_ is the volume of the red blood cells; *Q*_1_(*x*) denotes the flow of the fluid filtration/refilling between the two-compartments across the capillary membrane, which is governed by the Starling’s equation [11]; *Q*_2_(*x*_2_) describes the lymphatic flow, we have

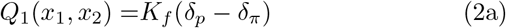

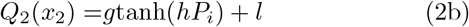

where *δ*_*p*_ and *δ*_*π*_ represent the hydrostatic and osmotic pressure gradients, respectively. The mathematical descriptions of these functions are given in the Appendix Appendix A. *g, h*, and *l* are constants listed in Table A.1. During the fluid exchange between the two compartments, *Q*_2_(*x*_2_) is smaller that *Q*_1_(*x*) [27].

## 3. Nonlinear Stability Analysis

The class of nonlinear compartment models that we are dealing with in this paper is developed in [22, 24, 25]. In order to derive a general stability result (Theorem 1 below) for this class, we consider *n* compartments where the dynamics of a certain amount of a material of interest in the *i* th compartment at time *t*, denoted by *x*_*i*_(*t*), is given by

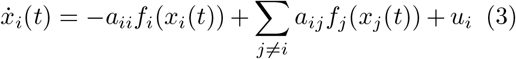

where *a*_*ij*_ ≥ 0, *i, j* ∈ {1, …, *n*}. The *i*th compartment receives the material through the input at rate *u* and from some other neighboring compartments (e.g., *j*th) at rate *a*_*ij*_*f*_*j*_(*x*_*j*_(*t*)) where the nonlinear function *f*_*j*_(*x*_*j*_) depends on the concentration of the *j*th compartment *x*_*j*_.

Note that all the results of stability in these studied [22, 24, 25] were based on the assumption that *f*_*ij*_(*x*_*j*_) ≥ 0, for *x*_*j*_ ≥ 0, 1≤ *i, j*≤ *n, i* ≠ *j* and *f*_*ij*_(0) = 0. However, these results do not imply to the model in (1) because *Q*_1_(*x*) ≤ 0 (non-positive) and the nonlinear functions *Q*_1_(*x*) and *Q*_2_(*x*_2_) defined in (2a) and (2b), respectively, are not continuous in the entire domain of *x* ∈ ℝ^2^. This readily can be seen from (A.4), in words at the origin (*x* ≡ 0). The dynamics in (3) with *n* compartments has a more compact form, which can be written as

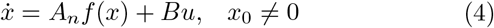

where *f* (*x*) = [*f*_1_(*x*_1_), …, *f*_*n*_(*x*_*n*_)]^*T*^, *B* is an *I*_*n*_ identity matrix, and the matrix *A*_*n*_ ∈ ℝ^*n×n*^ is a symmetric matrix (i.e., *a*_*ij*_ = *a*_*ji*_) defined by

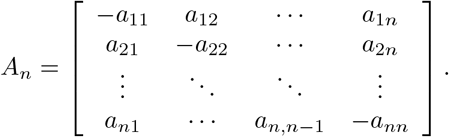

Note that the nonlinear function of the flow *Q*_1_(*x*_1_, *x*_2_) (2a) depends on both states *x*_1_ and *x*_2_, whereas, in the general form (3), the flow of the compartment *i*, (i.e., *f*_*i*_(*x*_*i*_)), depends only on the associated state *x*_*i*_. We illustrated the rearrangement of *Q*_1_ in the form of (3) in the next section.

### 3.1. Compact Form and Equilibrium Set

It is possible to rearrange the model (1) in the compact form of (4) by rearranging the equation of *Q*_1_(*x*) with *Q*_2_(*x*_2_). This rearrangement is explicitly derived in the Appendix in Appendix B, (B.1). The compact form of the model in (B.1) is:

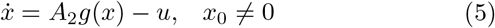

where *g*(*x*) = [*g*_1_(*x*_1_) *g*_2_(*x*_2_)]^*T*^ and *A*_2_ is symmetric, negative semi-definite given by

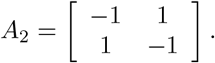

The mathematical description of *g*_1_(*x*_1_) and *g*_2_(*x*_2_) are detailed in Appendix Appendix B, (B.1). Because both of these functions are not continuous in the entire domain of *x* ∈ ℝ^2^, we specify a restrictive domain as follows by this subset:

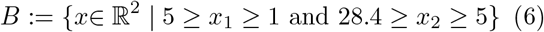

This closed subset is started at (1, 5) liters and closed at (5, 28.4) liters, which is illustrated by Fig.2. A brief description is provided in the Appendix Appendix A about how we specified this subset. Note that since the model (5) is defined only in the set *B* (6), an unstable solution of (5) refers to a solution *x*(*t*) (e.g., at *t* ≡ 0, *x* ∈ *B*) leaves the set *B* as *t* → ∞.

**Figure 2:**
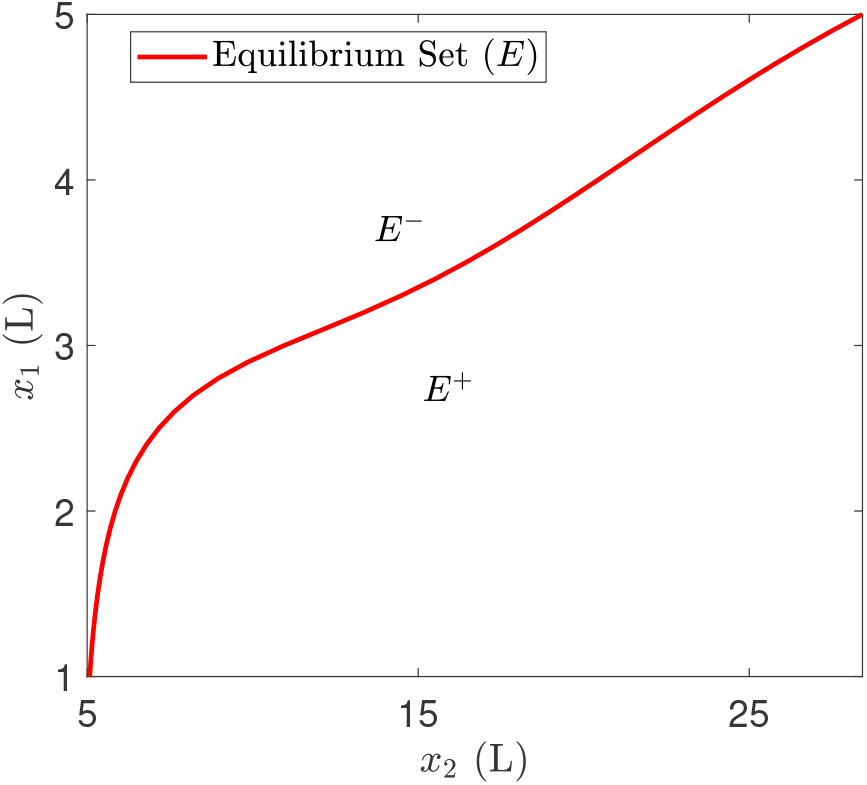
The equilibrium set *E* of the model (5) for all *x* ∈ *B*, including *E*^+^ and *E*^*−*^ subsets. *E* starts at an equilibrium point (1, 5) and ends at this equilibrium point (5, 28.4).

From a mathematical point of view, fluid volume homeostasis can be seen as a stable equilibrium point. To obtain an equilibrium point (*x*_*eq*_), we assume that there is no UFR (*u* ≡ 0), then we solve for *g*_1_(*x*_1*eq*_) = *g*_2_(*x*_2*eq*_) where this equality occurs at equilibrium: 0 = *A*_2_*g*(*x*_*eq*_). Due to high nonlinearity in these functions: *g*_1_(*x*_1_) and *g*_2_(*x*_2_), we could not derive an explicit expression, analytically, for the equilibrium point, *x*_*eq*_, however, we compute instead an equilibrium set *E*, numerically, as shown in Fig. 2, by defining *x*_1_ and *x*_2_ which satisfy the following relation: *g*_1_(*x*_1_) = *g*_2_(*x*_2_).

We define the equilibrium set for the model in (5) as follows:

#### Definition 1.

*Consider model (5) with u* ≡ 0, *the equilibrium set of (5) is define by*

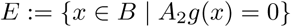

*in which g*_1_(*x*_1_) = *g*_2_(*x*_2_).

By defining the net flow as: *G*(*x*) = −*g*_1_(*x*_1_) + *g*_2_(*x*_2_), we can introduce two subsets above and below *E*, described by *E*^+^ ≔ {*x* ∈ *B* | *G*(*x*) *>* 0} and *E*^*−*^ ≔ {*x* ∈ *B* | *G*(*x*) *<* 0}, as shown in Fig. 2.

## 4. Results

Our stability results in this section is discussed as follows. First, we deriving new global stability results for the general class of nonlinear model (4) using an appropriate Lyapunov function. This model consists of *n* compartments and with zero input (*u* ≡ 0), i.e., 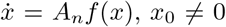. Then we apply this result to our model in (5).

Second, under nonzero input *u* ≠ 0 (UFR) and certain physiological conditions, we demonstrate the boundedness of the solution *x*(*t*) of (5) during 0 ≤ *t <* ∞. Specifically, we use qualitative properties of *G*(*x*) and under various forms of UFR profiles used in dialysis therapy to ensure that the solution *x*(*t*) of (5) is contained in a closed pre-defined set during *t* ≥ 0 and then as *t* → ∞, *x*(*t*) approaches the equilibrium set *E*. Inside this closed set, we explore the behavior of *x*(*t*) to gain physiological insights into fluid dynamics.

The fluid volume in each compartment of our model has a dynamics defined in (5). This dynamics can be determined by a particular slope *s*. When *u* ≡ 0 in (5), the slope is given very simply as follows:

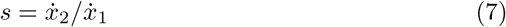

### Lemma 1.

*When u*_1_ ≡ 0 *in (5), the slope s* = −1 *for all x* ∈ *B* − *E*.

*Proof*. We substitute 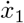 and 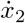 from (5) into (7), thus

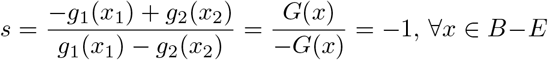

□

Note that for all points in *E*, by definition 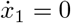, 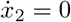 consequently, the slope is undefined.

### 4.1 Global Stability Results

We will establish the global stability result of the general compartment model in (4). For this purpose, we invoke LaSalle’s Theorem since it applies to a system having an equilibrium set rather than an isolated equilibrium point [28]. The main results of this paper are demonstrated in Theorem 1 and Theorem 2, by which nonlinear stability of a class of compartment models (4) and the boundedness of the solution of (5) are explored.

#### Theorem 1.

*Consider (4) with u* ≡ 0 *and let* Ω ⊂ ℝ^*n*^ *be compact and positively invariant with respect to the system in (4). Assume f* (*x*) *is differentiable in* Ω, *and let E be the equilibrium set E* = {*x* ∈ Ω | *A*_*n*_*f* (*x*) = 0}, *and let* 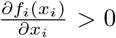, ∀*x* ∈ Ω. *If x*_0_ ∈ Ω, *then x*(*t*) *approaches E as t* → ∞.

*Proof*. Consider the Lyapunov function candidate: *V* (*x*) : ℝ^*n*^ → ℝ defined by

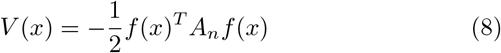

where this function is continuous, differentiable. Note *f* (*x*) and *A*_*n*_ are given in (4). The derivative of *V* (*x*) with respect to the time is derived in the Appendix Appendix C as

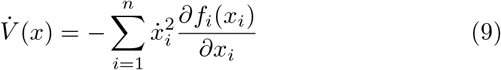

where 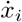 is defined in (3). This function satisfies the following conditions:

- 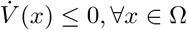,
- 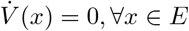.

In fact, *E* is the largest invariant subset in which 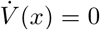, this can be shown as follows. Obviously *E* is a subset of 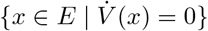. Now suppose *x* is such that 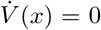. Then by (9) 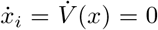 for all *i*, therefore, *x* ∈ *E*. It follow by LaSalle’s Theorem (Theorem 4.4 in [29]), that *x*(*t*) converges to *E*. □

In the next subsection, we will invoke Theorem 1 to prove global stability of the fluid volume two-compartment model.

### 4.2. Application to Two-compartment Model

We will demonstrate the attractivity of the equilibrium set of the two-compartment model which defined in (5). To invoke Theorem 1, we consider model (4) with *n* = 2. For this model, the 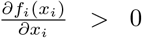, *i* = 1, 2, ∀*x* ∈ *B* for the parameters given in Table A.1. Consider the Lyapunov function:

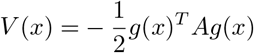

where in this case, matrix *A* is as defined in (5). Then

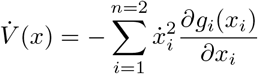

where 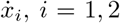 are given in (1). We simplify 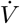 as

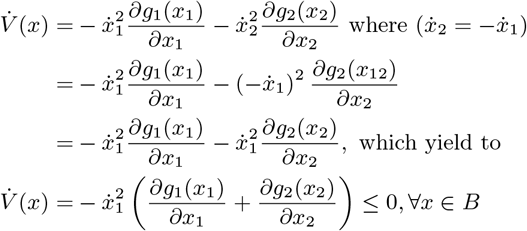

By construction the subset *B* is a compact set. To show that *B* is a positively invariant set with respect to model (5), from (Lemma 1) if *x*_0_ ∈ *B*, then *x*(*t*) approaches *E* as *t* → ∞ with a particular slope and direction such that *x*(*t*) never leave *B*.

### 4.3. Boundedness of Solutions with Nonzero Input

To examine the boundedness of the solution of under nonzero input (*u* ≠ 0), we use qualitative properties of *G*(*x*) under the effect of the UFR profile.

For this purpose, we derive the partial derivatives 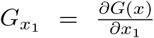 and 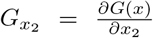 (see Appendix Appendix C.1) where *G*(*x*) is defined previously as *G*(*x*) = −*g*_1_(*x*_1_) + *g*_2_(*x*_2_). We also define the target volume to be removed as *V*_*T*_ (L), and 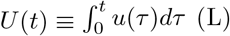 is the total amount of fluid volume removed at time *t* determined by accumulating UFR (*u*) during [0, *t*]; therefore, the removed fluid volume at the end of a dialysis session is 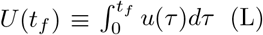 where *t*_*f*_ is the duration of the dialysis session; *V* (*t*) = *x*_1_(*t*) + *x*_2_(*t*) (L) is defined as the total fluid volume in the system at time *t*; and *V*_0_ = *x*_10_ + *x*_20_ (L) is the initial fluid volume in the system. Theorem 2 demonstrates that the solution *x*(*t*) of (5) is stable and contained in a per-defined closed set: *B*_*m*_ ≔ {*x* : 0 ≤ *G*(*x*) ≤ *u*_*max*_, *x*_1_ ≤ *x*_10_, *x*_2_ ≤ *x*_20_, and (*x*_1_ + *x*_2_) *> V*_0_ − *U* (*t*_*f*_). In this set, we use physiological values (from dialysis therapy) for all these parameters: UFR profile, *u*_*max*_, *x*_10_, *x*_20_, and *t*_*f*_ to examine the boundedness of the solution *x*(*t*) for *t* ≥ 0 and then as *t* → ∞ (i.e., *t > t*_*f*_), thus *x*(*t*) approaches *E*. However, as we highlighted earlier, since the model (5) is defined only in the set *B* (6), an unstable solution of (5) means that *x*(*t*) leaves the set *B*.

#### Theorem 2.

*Consider (5) with x*_0_ ∈ *E where E* ≔ {*x* ∈ *B* : *G*(*x*) = 0}. *Let E*^+^ ≔ {*x* : *G*(*x*) *>* 0}. *Suppose G is continuous on the interior of the 1st quadrant and satisfies the following:*

1. 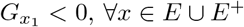.
2. 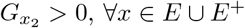.

A. *For given:* 0 *< u*_*max*_ *< U*_0_, *V*_*T*_ ≤ *x*_10_, *u*(*t*) ≥ 0, *and t*_*f*_ *>* 0, *take input u*_1_(*t*):

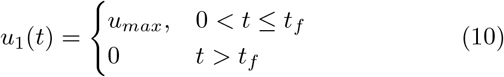
B. *For given:* 0 *< u*_*max*_ *< U*_0_, *V*_*T*_ ≤ *x*_10_, *û* (*t*) ≥ 0, *and t*_*f*_ *>* 0, *take input u*_2_(*t*):

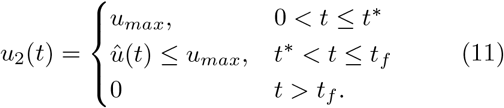

*Let x*(*t*) *be a solution to (5) and let B*_*m*_ *be: B*_*m*_ ≔ {*x* : 0 ≤ *G*(*x*) ≤ *u*_*max*_, *x*_1_ ≤ *x*_10_, *x*_2_ ≤ *x*_20_, *and* (*x*_1_ + *x*_2_) *> V*_0_ − *U* (*t*_*f*_)}. *Then x*(*t*) ∈ *B*_*m*_ *for all* 0 ≤ *t* ≤ *t*_*f*_.

The prove of Theorem 2 is provided in the Appendix Appendix D.

We will illustrate this result by considering the following numerical examples.

## 5. Simulation Examples

To test our results of the boundedness of the solution under a nonzero input, we consider the model in (5) with different UFR profiles (see Fig. 3) applied to the system, including constant UFR profile, step UFR profile, and patient-specific (time-varying) UFR profile developed in our previous work [30], as shown in Fig. 3, bottom. This UFR profile was designed to meet the decrease in fluid refilling during the last two-thirds of the therapy.

**Figure 3:**
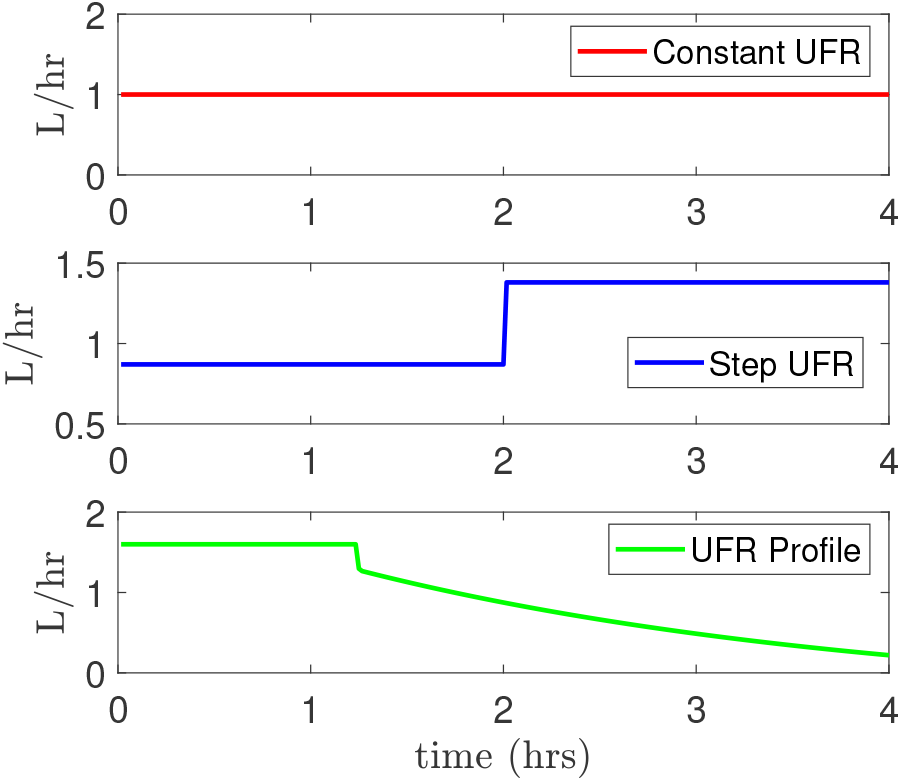
Top: A constant UFR profile, Middle: Step UFR profile, and Bottom: patient-specific (time-varying) profile used to explore the boundedness and behavior of the solution *x* during 0 ≤ *t < t*_*f*_.

In Theorem 2, we give physiological conditions under which the solution *x*(*t*) is contained in this set *B*_*m*_ where *B*_*m*_ ≔ {*x* : 0 ≤ *G*(*x*) ≤ *u*_*max*_, *x*_1_ ≤ *x*_10_, *x*_2_ ≤ *x*_20_, and (*x*_1_ +*x*_2_) *> V*_0_ −*U* (*t*_*f*_)}. During 0 ≤ *t < t*_*f*_, *x*(*t*) will be bounded in this region: 0 ≤ *G*(*x*) ≤ *u*_*max*_ where *u*_*max*_, which describes the maximum UFR rate in clinical practices should be *u*_*max*_ ≤ 13 ml/hr/kg to minimize cardiovascular risk [31]. As shown in Fig. 3, we chose the duration of the dialysis treatment to be *t*_*f*_ = 4 hrs. From the UFR profiles in Fig 3, the target fluid volume (*V*_*T*_) was as follows: 4L, 4.5L, and 4L, respectively. The initial conditions (initial volumes) for these dialysis sessions were determined to satisfy this condition: *V*_*T*_ ≤ *x*_10_ (Theorem 2). This condition allows to begin the dialysis therapy with an initial fluid volume *V*_0_ = *x*_10_ + *x*_20_ that is a large enough to keep *x*(*t*) inside the set *B*_*m*_ during 0 ≤ *t < t*_*f*_. As we illustrated in Fig. 4 and 6, since *V*_*T*_ = 4L, we used initial conditions (4, 20.4)L located on *E*, whereas, for the step UFR profile (see Fig. 5), the initial conditions were at (4.5, 24.2)L because the target volume was *V*_*T*_ = 4.5L.

**Figure 4:**
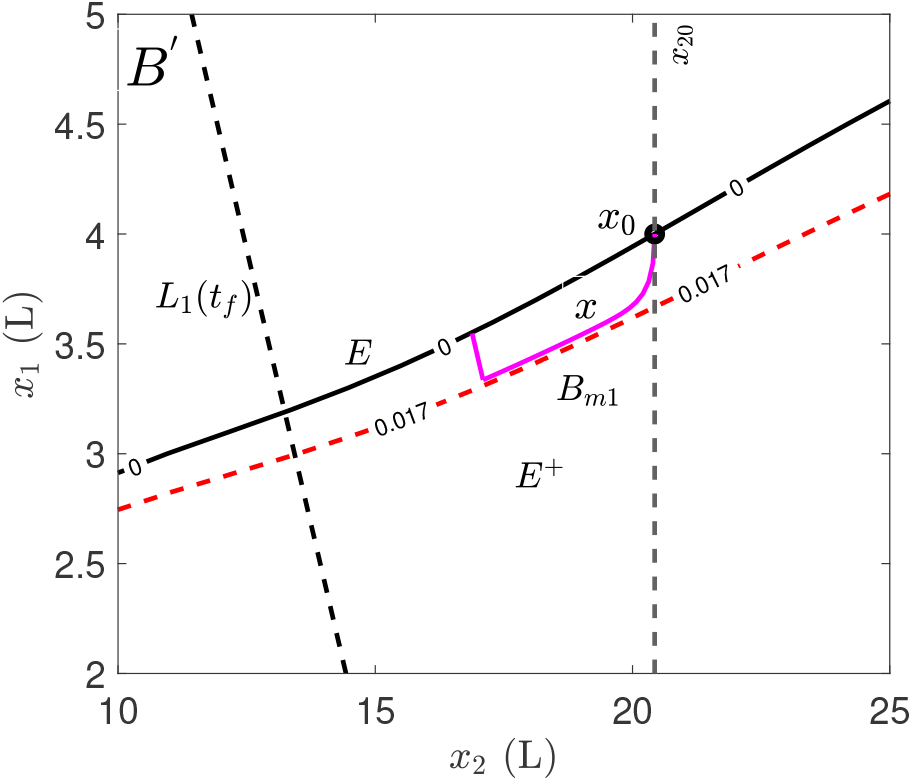
The solution *x*(*t*) was contained in the set *B*_*m*1_ under the constant UFR profile. The set *B*_*m*1_ is determined as: *B*_*m*1_ ≔ {*x* : 0 ≤ *G*(*x*) ≤ 0.017 [L/min], *x*_1_ ≤ 4L, *x*_2_ ≤ 20.4L, and *L*_1_(*t*_*f*_) = (*x*_1_ + *x*_2_) *>* 20.4L}, where *B′* ⊂ *B*.

**Figure 5:**
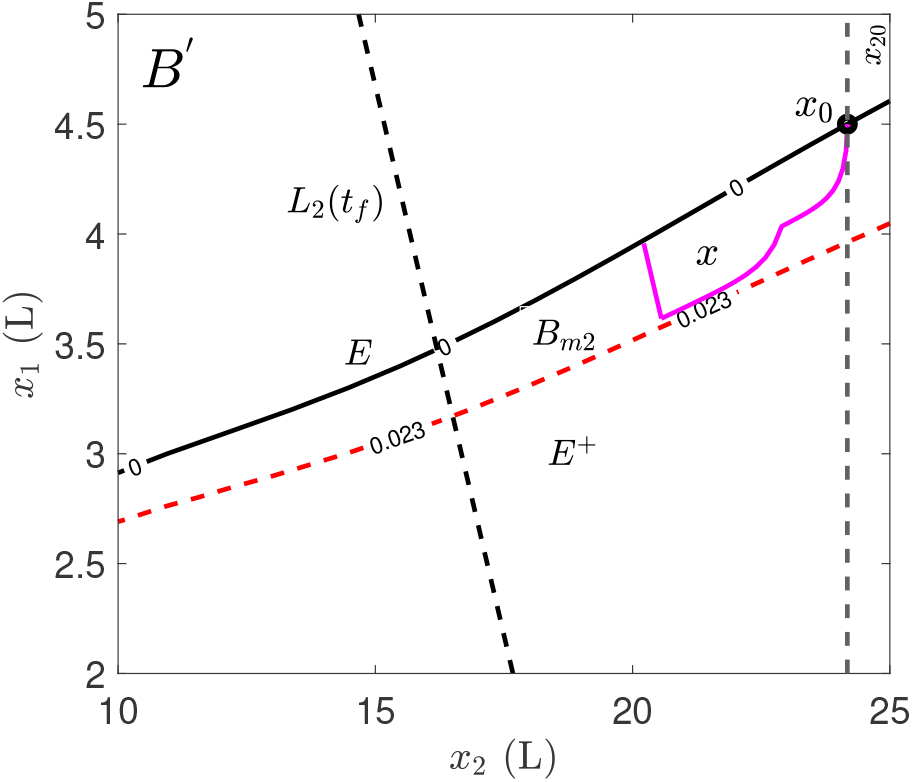
The solution *x*(*t*) was contained in the set *B*_*m*2_ under the step UFR profile. The set *B*_*m*1_ ≔ {*x* : 0 ≤ *G*(*x*) ≤ 0.023 [L/min], *x*_1_ ≤ 4.5L, *x*_2_ ≤ 24.2L, and *L*_1_(*t*_*f*_) = (*x*_1_ + *x*_2_) *>* 24.2L}, where *B*^*′*^ ⊂ *B*.

**Figure 6:**
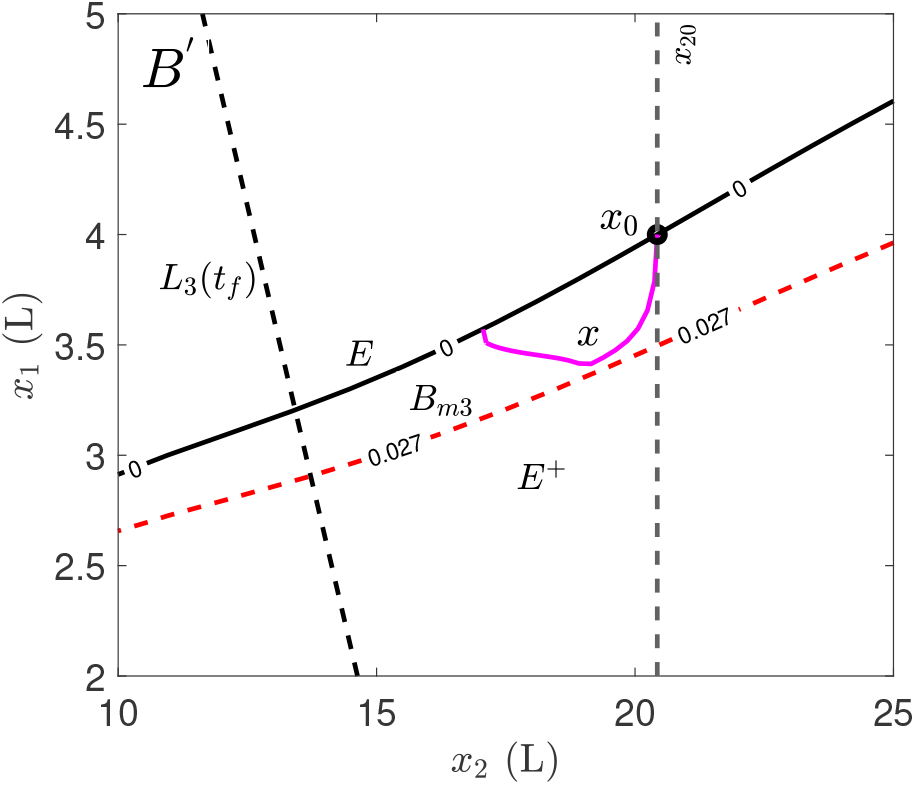
The solution *x* was contained in the set *B*_*m*3_ under the time-varying UFR profile. The *B*_*m*3_ is determined by *B*_*m*3_ ≔ {*x* : 0 ≤ *G*(*x*) ≤ 0.027[L/min], *x*_1_ ≤ 4L, *x*_2_ ≤ 20.4L, and *L*_3_(*t*_*f*_) = (*x*_1_ + *x*_2_) *>* 20.6L}, where *B*^*′*^ ⊂ *B*.

The time-varying UFR profile (Fig. 3, bottom) was designed to remove more fluid in the first one-third of the session (0.027L/min, which is 1.62 L/hr) when the fluid refilling rate is high, and then the UFR was decreased slightly towards the end of the session when the refilling rate could be low to prevent intradialytic hypotension. However, the step UFR profile (Fig. 3, middle) was increased in the second half of the therapy, which is not recommended in clinical practices because it may provoke intradialytic hypotension due to a significant reduction in the blood volume.

We illustrated the boundedness of the solution *x*(*t*) under the three UFR profiles by the Phase-plane in Fig. 4, 5, and 6. The solution *x*(*t*) of (5) was contained in these sets during the time frame 0 *< t* ≤ *t*_*f*_, respectively:

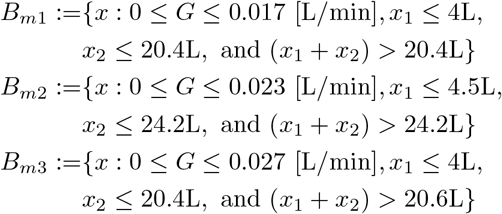

*V*_*T*_ ≤ *x*_10_ To provide physiological insights and a deep understanding of fluid removal during the dialysis treatment, we explore the behavior of the solution *x*(*t*) inside these sets. Inside *B*_*m*1_ and *B*_*m*2_, from the solution’s trajectory *x*(*t*), there was a reduction in the fluid volume in both directions (*x*_1_, *x*_2_) under the constant and step function UFR profiles during the entire session 0 *< t* ≤ *t*_*f*_. A significant reduction in *x*_1_ provokes intradialytic hypotension. The reduction in *x*_1_ was significantly appeared under the step function UFR profile (Fig. 5), due to the step increased in the UFR. In addition, from a mathematical perspective, with constant and step function UFR profiles, *x*(*t*) may leave the set *B* and becomes unstable if *x*_10_ started in the left lower corner of *B* set, which is closer to the lowest value 1L. This case can be seen when the condition *V*_*T*_ ≤ *x*_10_ is violated. On the other hand, the time-varying UFR profile, which removes the same amount of fluid (4L), has an opposite effect on the trajectory *x*(*t*) compared with the constant UFR profile. The trajectory *x*(*t*) was considerably increased (*x*_1_ increased) in the second half of the session, as shown in Fig. 6. This increase in *x*_1_ could prevent the occurrence of intradialytic hypotension, which likely occurs at the end of each session.

### 5.1. Discussions

We illustrated a global stability analysis applied to a specific class of nonlinear models used in dialysis treatment. We used qualitative properties of the net flow *G*(*x*) and UFR profiles to ensure the boundedness of the solution of our model. Our results indicated that if the dialysis session started in the physiological set *B*_*m*_, the model’s solution never leaves this set.

To explore the solution’s behavior under ultrafiltration, we compared the standard UFR profiles’ effect versus the patient-specific profile on the solution’s trajectory *x*(*t*) inside *B*_*m*_ during the dialysis therapy. The slandered UFR profiles do not guarantee optimal dialysis outcomes. In contrast, the time-varying UFR profile, which was designed based on physiological insights to meet the lack in vascular refilling rate, could lead to effective and safer dialysis treatment.

## 6. Conclusions

The current focus on improving fluid management in dialysis therapy motivated the need for exploring stability analysis for a specific class of models used in artificial kidney treatment in end-stage kidney disease patients.

This paper presented a novel global stability analysis for a class of nonlinear multi-compartment models whose flow functions are non-positive. These results were applied to our nonlinear two-compartment model. We demonstrated the solution’s boundedness and behavior of the two-compartment model during dialysis therapy to gain physiological insights into fluid removal risk. We showed that under physiological conditions, the nonlinear model’s solution could be bounded and contained in a pre-defined set during the ultrafiltration. We concluded that the solution’s trajectory under the patient-specific UFR profile has better performance than the standard ultrafiltration profiles. These results motivated us to provide new and more advanced therapeutics for dialysis treatment, to save numerous lives of people with end-stage kidney disease.

## Acknowledgement

The authors wish to acknowledge the contribution of Dr. C.V.Hollot in setting and proving the theorems.

## Appendix A. Model Description

From (2) we have

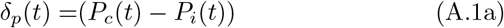

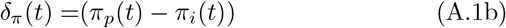

where *Pc, Pi, π*_*p*,_ and *π*_*i*_ refer to the hydrostatic capillary pressure, interstitial pressure, plasma colloid osmotic pressure, and interstitial colloid osmotic pressure, respectively. The capillary pressure is defined by

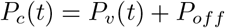

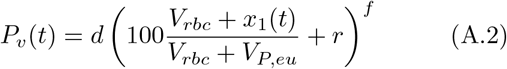

where *P*_*v*_ is the venous pressure, *P*_*off*_ is a pressure offset, *x*_1_(*t*) is the intravascular volume at a given time *t, V*_*P,eu*_ is intravascular volume at the euhydrated state, *d, e* and *l* are constants [32] listed in Table A.1. We define

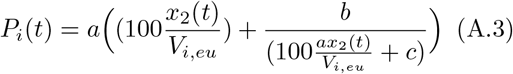

where *x*_2_(*t*) represents interstitial fluid volume at a given time *t, V*_*i,eu*_ refers to interstitial volume at the euhydrated state, *a, b* and *c* are constants [32], and where

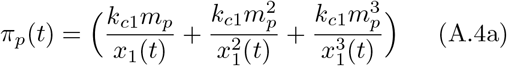

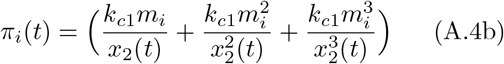

where *m*_*p*_ and *m*_*i*_ describe the protein mass in plasma and interstitial volumes, respectively, *k*_1_, *k*_2_ and *k*_3_ are constants [33] (see Table A.1).

Since *x*_1_ and *x*_2_ represent volume, they should be positive. Hence, we will restrict the domain of *x*_1_ and *x*_2_ over this range *x*_1_ *>* 0, *x*_2_ *>* 0. Therefore, we set the term inside the bracket (·) in (A.2) to be greater than zero, then we solve for *x*_1_ accordingly. We repeat this step for the denominator in (A.3) to solve for *x*_2_. Note that the constants in (Table A.1) are used here. As a result, *Q*_1_(*x*) and *Q*_2_(*x*) are continuous in the following restricted domain:*x*_1_ ≥ 1 and *x*_2_ ≥ 5.

**Table A.1:** Model parameters

| Para. | Value | Para.-Unit | Value |
| --- | --- | --- | --- |
| a | 0.0065 | $K_f$ [L/min*mmHg] | 0.0052 |
| b | -146.3 | $m_p$[g] | 210 |
| c | -54.72 | $m_i$[g] | 210 |
| d | 0.01 | $P_{off}$ [mmHg] | 13 |
| r | -30.0 | $k_1$ | 0.21 |
| f | 1.5 | $k_2$ | 0.0016 |
| g | 0.02 | $k_3$ | 9e-6 |
| h | 0.63 | $V_{P,eu}$ (L) | 3 |
| l | 0.02 | $V_{i,eu}$ (L) | 12 |
| - | - | $V_{rbc}$ (L) | 2 |

## Appendix B. Compartmental Model Arrangement

We rearrange *Q*_1_(*x*) by separating the terms that contain *x*_1_ and *x*_2_ as follows: *Q*_1_(*x*_1_, *x*_2_) = *K*_*f*_ (*P*_*c*_ − *π*_*p*_) − *K*_*f*_ (*P*_*i*_ − *π*_*i*_) and then substitute the later *Q*_1_(*x*) in the model (3):

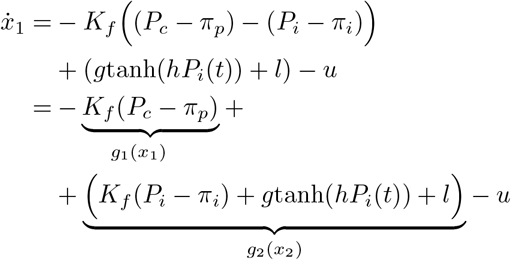

and 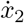 becomes:

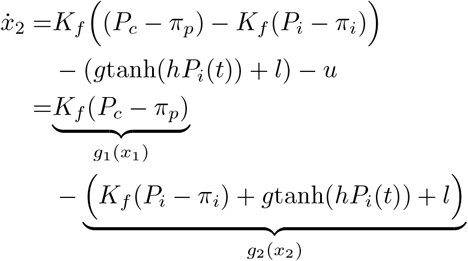

that leads to

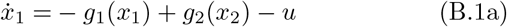

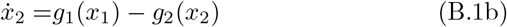

where *g*_1_*(x*_1_*) = K*_*f*_ *(P*_*c*_*− π*_*p*_) and *g*_2_(*x*_2_) = *K*_*f*_ (*P*_i_ −*π*_i_) + *gtanh*(*hP*_i_(*t*)) + *l*.

## Appendix C. Theorem 1

*Proof*. We have this Lyapunov candidate for *n* compartments:

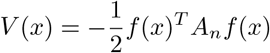

Substituting *f* ^*T*^ (*x*) = [*f*_1_(*x*_1_) … *f*_*n*_(*x*_*n*_)] and *A*_*n*_ (4) into *V* (*x*) leads to:

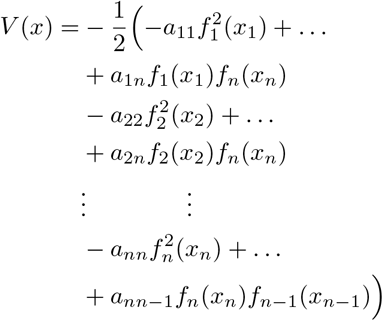

The derivative 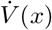 is:

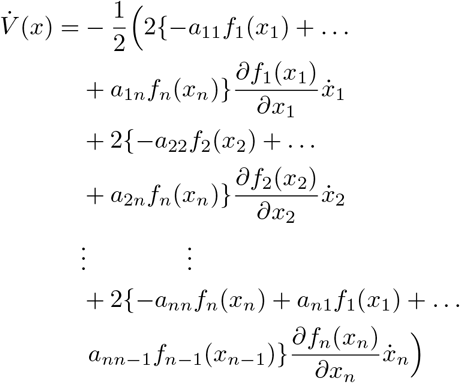

This gives:

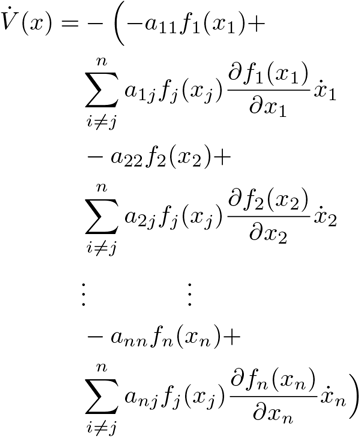

From (3), we have the following:

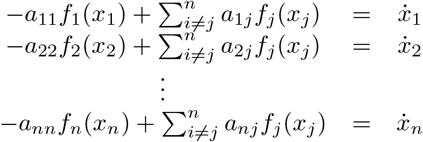

Thus

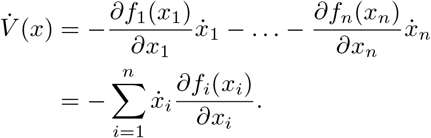

### Appendix C.1. Partial Derivative of the Flow

The partial derivative terms: 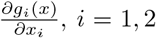 are derived here:

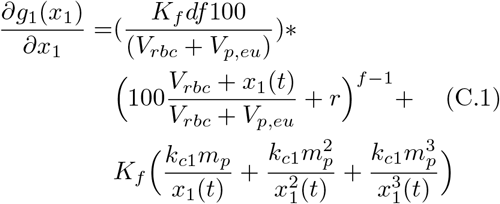

and

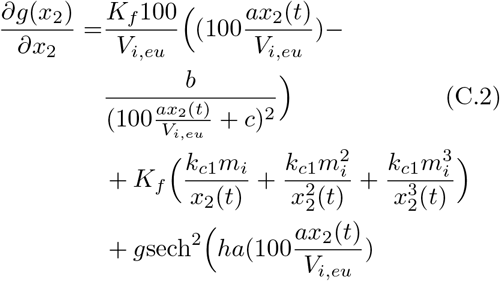

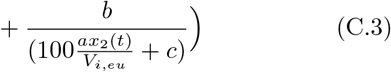

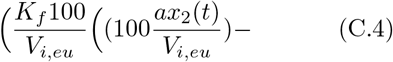

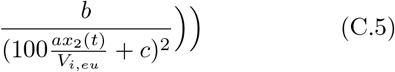

Using the parameters in Table A.1, we have: 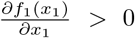 and 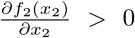. Note that since *G*(*x*) = −*g*_1_(*x*_1_) + *g*_2_(*x*_2_), then 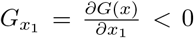 and 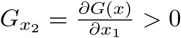. □

## Appendix D. Theorem2 (proof)

Substituting *G*(*x*) = − *g*_1_(*x*_1_) + *g*_2_(*x*_2_) into (5) leads to

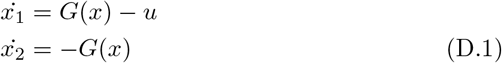

and then

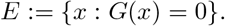

*Proof*. Along the solution to (5), 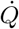 is given by

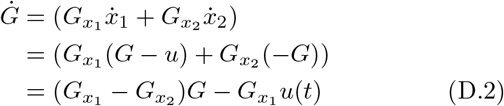

a. At *G*(*x*) = 0, from (D.2) and condition (1), 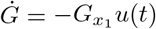 :

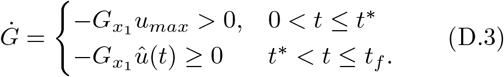
b. Using *G*(*x*) = *u*_*max*_, from (D.2) and condition (2) 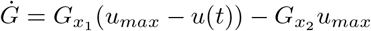:

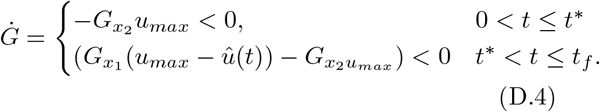
c. Using *G*(*x*) ≤ *u*_*max*_, it follows:

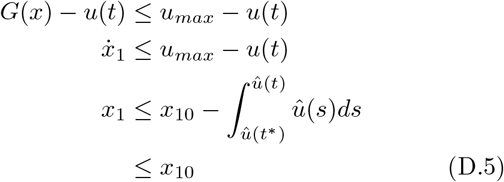
d. and using *G*(*x*) ≥ 0, it follows:

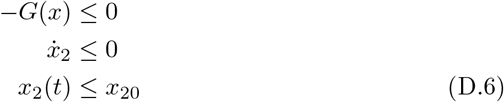

From (5), we have

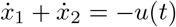

then

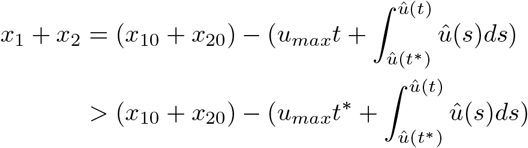

then from a, b, and d: *x*(*t*) ∈ *B*_*m*_, for all 0 ≤ *t* ≤ *t*_*f*_. This proves the Theorem. □

## References

[1] Jennifer E Flythe, Matthew J Tugman, Julia H Narendra, Magdalene M Assimon, Quefeng Li, Yueting Wang, Steven M Brunelli, and Alan L Hinderliter. Effect of ultrafiltration profiling on outcomes among maintenance hemodialysis patients: a pilot randomized crossover trial. Journal of Nephrology, pages 1–11, 2020.

[2] Susumu Ookawara, Kiyonori Ito, Takayuki Uchida, Keito Tokuyama, Satoshi Kiryu, Takeshi Suganuma, Kyoko Hojyo, Haruhisa Miyazawa, Yuichiro Ueda, Chiharu Ito, et al. Hemodialysis crossover study using a relative blood volume change-guided ultrafiltration control compared with standard hemodialysis: the bvufc study. Renal Replacement Therapy, 6(1):1–10, 2020.

[3] Jack Q Jaeger and Ravindra L Mehta. Assessment of dry weight in hemodialysis: an overview. Journal of the American Society of Nephrology, 10(2):392–403, 1999.

[4] Stefano Marano, Marco Marano, and Leandro Pecchia. Frontiers in hemodialysis part ii: Toward personalized and optimized therapy. Biomedical Signal Processing and Control, 61:102029, 2020.

[5] John A Sargent, Marco Marano, Stefano Marano, and F John Gennari. Acid-base homeostasis during hemodialysis: New insights into the mystery of bicarbonate disappearance during treatment. In Seminars in dialysis, volume 31, pages 468–478. Wiley Online Library, 2018.

[6] Jing Lin, Zhen Cheng, Xiaoqiang Ding, and Qi Qian. Acid-base and electrolyte managements in chronic kidney disease and end-stage renal disease: Case-based discussion. Blood purification, 45(1-3):179–186, 2018.

[7] Donal N Reddan, Lynda Anne Szczech, Vic Hasselblad, Edmund G Lowrie, Robert M Lindsay, Jonathan Himmelfarb, Robert D Toto, John Stivelman, James F Winchester, Linda A Zillman, et al. Intradialytic blood volume monitoring in ambulatory hemodialysis patients: a randomized trial. Journal of the American Society of Nephrology, 16(7):2162–2169, 2005.

[8] Biff F Palmer and William L Henrich. Recent advances in the prevention and management of intradialytic hypotension. Journal of the American Society of Nephrology, 19(1):8–11, 2008.

[9] Patrick B Reeves and Finnian R Mc Causland. Mechanisms, clinical implications, and treatment of intradialytic hypotension. Clinical journal of the American Society of Nephrology, 13(8):1297–1303, 2018.

[10] Rammah Abohtyra, CV Hollot, Michael G Germain, Yossi Chait, and Joseph Horowitz. Personalized ultrafiltration profiles to minimize intradialytic hypotension in end-stage renal disease. In 2018 IEEE Conference on Decision and Control (CDC), pages 309–314. IEEE, 2018.

[11] A Aurelio, Doris H Fuertinger, Franz Kappel, Anna Meyring-Wõsten, Stephan Thijssen, and Peter Kotanko. A physiologically based model of vascular refilling during ultrafiltration in hemodialysis. Journal of theoretical biology, 390:146–155, 2016.

[12] John T Daugirdas. Pathophysiology of dialysis hypotension: an update. American journal of kidney diseases, 38(4):S11–S17, 2001.

[13] Arthur C Guyton and JE Hall. The body fluid compartments: extracellular and intracellular fluids; interstitial fluid and edema. Textbook of medical physiology, 11:293, 2000.

[14] Jurgen Floege, Richard J Johnson, and John Feehally. Comprehensive clinical nephrology. Elsevier Health Sciences, 2010.

[15] John E Hall. Pocket companion to Guyton & Hall textbook of medical physiology. Elsevier Health Sciences, 2015.

[16] PW Chamney, C Johner, C Aldridge, M Kramer, N Valasco, JE Tattersall, T Aukaidey, R Gordon, and RN Greenwood. Fluid balance modelling in patients with kidney failure. Journal of medical engineering & technology, 23(2):45–52, 1999.

[17] Sarah Faubel and Joel Topf. The fluid, electrolyte & acid-base companion. Alert and Oriented Pub., 1999.

[18] PR Keshaviah, KM Ilstrup, and FL Shapiro. Dynamics of vascular refilling. Artif. Organs, 2:506–10, 1983.

[19] Daniel Schneditz, Johannes Roob, Martina Oswald, Helmuth Pogglitsch, Maximilian Moser, Thomas Kenner, and Ulrich Binswanger. Nature and rate of vascular refilling during hemodialysis and ultrafiltration. Kidney international, 42(6):1425–1433, 1992.

[20] Kaoru Tabei, Hirofumi Nagashima, Osamu Imura, Toshihiro Sakurai, and Yasushi Asano. An index of plasma refilling in hemodialysis patients. Nephron, 74(2):266–274, 1996.

[21] John A Jacquez and Carl P Simon. Qualitative theory of compartmental systems. Siam Review, 35(1):43–79, 1993.

[22] Reginald F Brown. Compartmental system analysis: State of the art. IEEE Transactions on Biomedical Engineering, (1):1–11, 1980.

[23] Wassim M Haddad, Tomohisa Hayakawa, and James M Bailey. Adaptive control for non-negative and compartmental dynamical systems with applications to general anesthesia. International journal of adaptive control and signal processing, 17(3):209–235, 2003.

[24] Hajime Maeda and Shinzo Kodama. Qualitative analysis of a class of nonlinear compartmental systems: nonoscillation and asymptotic stability. Mathematical Biosciences, 38(1):35–44, 1978.

[25] Hajime Maeda, Shinzo Kodama, and Yuzo Ohta. Asymptotic behavior of nonlinear compartmental systems: nonoscillation and stability. Circuits and Systems, IEEE Transactions on, 25(6):372–378, 1978.

[26] Airlie Chapman and Mehran Mesbahi. Stability analysis of nonlinear networks via m-matrix theory: Beyond linear consensus. In 2012 American Control Conference (ACC), pages 6626–6631. IEEE, 2012.

[27] AC Guyton and JE Hall. Textbook of medical physiology (9th edn) wb saunders company. Philadelphia. Pennsylvania, page 1006, 1996.

[28] Hassan K Khalil and JW Grizzle. Nonlinear systems, volume 3. Prentice hall New Jersey, 1996.

[29] Mathukumalli Vidyasagar. Nonlinear systems analysis, volume 42. Siam, 2002.

[30] Rammah Abohtyra, Yossi Chait, Michael J Germain, Christopher V Hollot, and Joseph Horowitz. Individualization of ultrafiltration in hemodialysis. IEEE Transactions on Biomedical Engineering, 66(8):2174–2181, 2018.

[31] John WM Agar. Personal viewpoint: Limiting maximum ultrafiltration rate as a potential new measure of dialysis adequacy. Hemodialysis International, 20(1):15–21, 2016.

[32] Aimee Stasia Goncalves. Model-based feedback control for hemodialysis treatment of chronic kidney disease patients, 2015.

[33] EM Landis and JR Pappenheimer. Exchange of substances through the capillary walls. Handbook of physiology, 2(2):961–1034, 1963.

